# Therapeutic liver cell transplantation to treat a genetic liver defect

**DOI:** 10.1101/2024.06.26.600783

**Authors:** Melanie Willimann, Hiu Man Grisch-Chan, Nicole Rimann, Tanja Rothgangl, Martina Hruzova, Gerald Schwank, Beat Thöny

**Author notes:** The first two authors contributed equally to this work. Corresponding author: Beat Thöny, University Children’s Hospital Zurich, Zurich Switzerland.

## Abstract

For gene therapy of the liver, *in vivo* applications based on adeno-associated virus are the most advanced vectors despite limitations, including low efficacy and episomal loss, potential integration and safety issues, and high production costs. Alternative vectors and/or delivery routes are of high interest. The regenerative ability of the liver bears the potential for *ex vivo* therapy using liver cell transplantation for disease correction if provided with a selective advantage to expand and replace the existing cell mass. Here we present such treatment of a mouse model of human of phenylketonuria (PKU). Primary hepatocytes from wild-type mice were gene modified *in vitro* (with a lentiviral vector) that carries a gene editing system (CRISPR) to inhibit *Cypor*. *Cypor* inactivation confers paracetamol (or acetaminophen) resistance to hepatocytes and thus a growth advantage to eliminate the pre-existing liver cells upon grafting (via the spleen) and exposure to repeated treatment with paracetamol. Grafting *Cypor*-inactivated wild-type hepatocytes into inbred young adult *enu2* (PKU) mice, followed by selective expansion by paracetamol dosing resulted in replacing up to 5% of cell mass, normalization of blood phenylalanine and permanent correction of PKU. Hepatocyte transplantation offers thus an armamentarium of novel therapy options for genetic liver defects.

## Introduction

Liver transplantation often is the last treatment options for patients suffering from severe inborn errors of metabolism (IEM). Unfortunately, the number of patients requiring liver transplantation surpasses its availability leaving many patients waiting for a suitable donor liver. Alternatively, liver gene therapy approaches or primary hepatocyte transplantation have been investigated as potential alternatives to liver transplantation theoretically, making the procedures safer, more accessible, and provide the potential for personalized and repeated treatment. Gene therapy is a broad field ranging from vaccinations to phenotype corrections for IEM^7-9^. Transfer vectors are indisputably a key component for successful gene delivery and safety, based on which adeno-associated viruses (AAVs) have emerged as most desirable delivery vectors. Over the years, multiple approaches have been developed to engineer AAV capsid and increase their specificity and transduction efficiency towards selected organs and tissues, in particular the liver^5^. Nevertheless, major restrictions for AAV-based approaches include manufacturing challenges^10^, limited packaging size, pre-existing immunity, induction of severe immune responses against viral vectors, and risks of insertional mutagenesis due to chromosomal integration^1-5,10-14^. Especially metabolic disorders pose an additional layer of complexity since IEM often require therapeutic intervention early in life where the liver is still growing rapidly, making episomal gene addition unsuitable due to loss of the therapeutic vector after cell replication^15^. Thus, modification of chromosomal DNA in hepatocytes is required for long-term treatment and targeted chromosomal integration of a therapeutic expression cassette, and transient expression of a prime or base editor provide ideal gene editing tools^16,17^.

The *Pah^enu2/enu2^* mouse is a model for human hyperphenylalaninemia or phenylketonuria (PKU)^18^ and has been extensively used in preclinical studies for liver gene therapy^19,20^. Both, gene addition and gene editing based approaches have been successful in phenotype correction where approximately 5-10% of hepatocytes must be targeted, independent of time point of administration^20,21^.

Even with highly promising gene editing tools for liver gene therapy, the limiting factor remains the delivery efficiency^15^. One approach to overcome limited gene targeting is to increase the fraction of targeted cells by *selection*. The liver is the ideal organ for selection due to its regenerative nature: damage or removal of parts of the liver will be compensated by repopulation through remaining hepatocytes. Hepatocyte transplantation has been investigated for decades due to its great potential but has not been successful until today due to the donor hepatocyte’s insufficient hepatic integrity and limited engrafting capacity^22^. Nevertheless, *in vivo* selective methods, based on *Fah* deficiency^23^, urokinase-type plasminogen activator (uPA)^24^, or most recently cytochrome P450 reductase (*Cypor*) inactivation,^6^ have been used successfully to expand genetically modified, allogenic, or xenogenic hepatocytes. Nonetheless, these methods require hepatocyte injury to host hepatocytes induced by toxic transgenes^25^ and relied on recipient mice being highly immunocompromised to decrease hepatocyte rejection and enable engraftment, such as the *Fah^-/-^, Rag2^-/-^, IL2rg^-/-^* (FRG) mouse. Recently, Tamaki *et al.* could show up to 70% stable liver repopulation upon autologous primary hepatocyte transplantation using fully immune competent mice, where allogenic primary hepatocyte transplantation was unsuccessful^26^. Unlike FAH deficient and uPA over-expressing hepatocytes, *Cypor*-inactivated hepatocytes have a selective advantage over “normal” hepatocytes making it a highly promising and versatile method to select for genetically modified hepatocytes. Paracetamol, also known as acetaminophen or APAP, is a well-studied drug used for temporary pain and fever relief. CYPOR is the main electron donor for cytochrome P450 enzymes (CYP) which metabolize paracetamol. It is metabolized in the pericentral regions of the liver and the majority is conjugated through glucuronidation (60%) and sulphation (30%) by uridine 5’-dishosho-glucoronosyltransferase (UGT) and sulfotransferase (SULT), respectively. Only 2-10% of paracetamol is oxidized by CYPs into the hepatotoxic metabolite N-acetyl-p-benzoquinone imine (NAPQI), which normally is immediately detoxified through glutathione conjugation^27,28^. In case of a paracetamol overdose, glucuronidation and sulfation pathways are saturated resulting in greater proportions of paracetamol being metabolized into NAPQI. Increased NAPQI quantities cannot be fully detoxified by glutathione stored in the liver, resulting in cell damage. Taking advantage of the paracetamol metabolism and its hepatotoxicity in case of a “mild” overdosing, hepatocytes can be genetically manipulated to become resistant to paracetamol by inactivating *Cypor*. Subsequently, CYP enzymes metabolizing paracetamol to NAPQI cannot be activated, thus hepatocytes become paracetamol resistant. It has been reported that *Cypor* inactivation and subsequent paracetamol-induced selection result in over 30-fold expansion and up to 50% replacement of the hepatic mass^6^. Therefore, by combining the delivery of a therapeutic gene with *Cypor* inactivation gene therapy approaches with insufficient gene delivery may still reach the therapeutic threshold and provide new avenues for cures for genetic liver defects.

In this work, we explored hepatocyte transplantation combined with subsequent *in vivo* paracetamol selection to treat the genetic metabolic disorder PKU. *Cypor*-inactivated wild-type donor hepatocytes were transplanted into PKU recipient mice, followed by graft-expansion using moderate paracetamol overdoses resulting in normalization of blood L-phenylalanine (Phe <360 μM) and correction of PKU. This work is the first proof of principle for an *ex vivo* gene therapy approach for the liver using primary hepatocytes (PH).

## Results

### *In vivo* testing of the gRNA-*SpCas9* gene editor to inactivate *Cypor*

To test the efficacy of a newly designed gRNA-*SpCas9* gene editor to inactivate *Cypor* in mouse liver, we used hydrodynamic tail vein (HTV) infusion of naked DNA that was reported to transfect up to 15% of hepatocytes^29-31^. Plasmid expression vector pLV.POR3-RFP (Fig. 1A; see also Methods section) was delivered into wild-type mice (n = 2; 100 μg of DNA equivalent to 7 x 10^12^ copies in each mouse). Indel frequency 8 weeks post-injection (and no paracetamol treatment) was below 2% by NGS (not shown). On the other hand, wild-type mice (n = 2) *not* injected with pLV.POR3-RFP but treated with 13 paracetamol doses showed indel frequency of up to 1.4% which was thereafter defined as background (Fig. 1B). Next, we performed HTV infusion of pLV.POR3-RFP (10 μg; 7 x 10^11^ copies) into wild-type mice (n = 5) and selected for *Cypor*-inactivated hepatocytes using paracetamol treatment or cycling. After 10 paracetamol doses, ALT responses remained between 50 and 800 U/l (for details see Methods). We thus analyzed the degree of indel frequency or hepatocyte selection in 3 mice after a total of 15 paracetamol doses and in 2 mice after 20 doses. As depicted in Fig. 1C-D, Immunohistochemistry (IHC) quantification after 15 doses revealed 34.6% *Cypor* inactivation while indel frequency using next generation sequencing (NGS) was 13.2%. In the 2 mice selected with 20 doses of paracetamol, we found no increase for indel frequency based on NGS analysis (11.2%). IHC analysis for CYPOR, e-Cadherin, and DAPI revealed, as expected, selective staining for pericentral (zone 3) and mid-lobular (zone 2) but not periportal hepatocytes (zone 1; Figure 1E-G). During establishing the paracetamol selection protocol which was based on Vonada *et al.*^6^, we noticed that extending the time between paracetamol doses from 3-4 days to 7 days was favorable in our hands as the ALT response increased. This implied that lower doses of paracetamol were required for moderate liver toxicity, and that more efficient and higher degrees of selection could be achieved with each dose (see Supplementary Fig. 1). For all the following experiments, mice were infused with one paracetamol dose per week.

**Figure 1.**
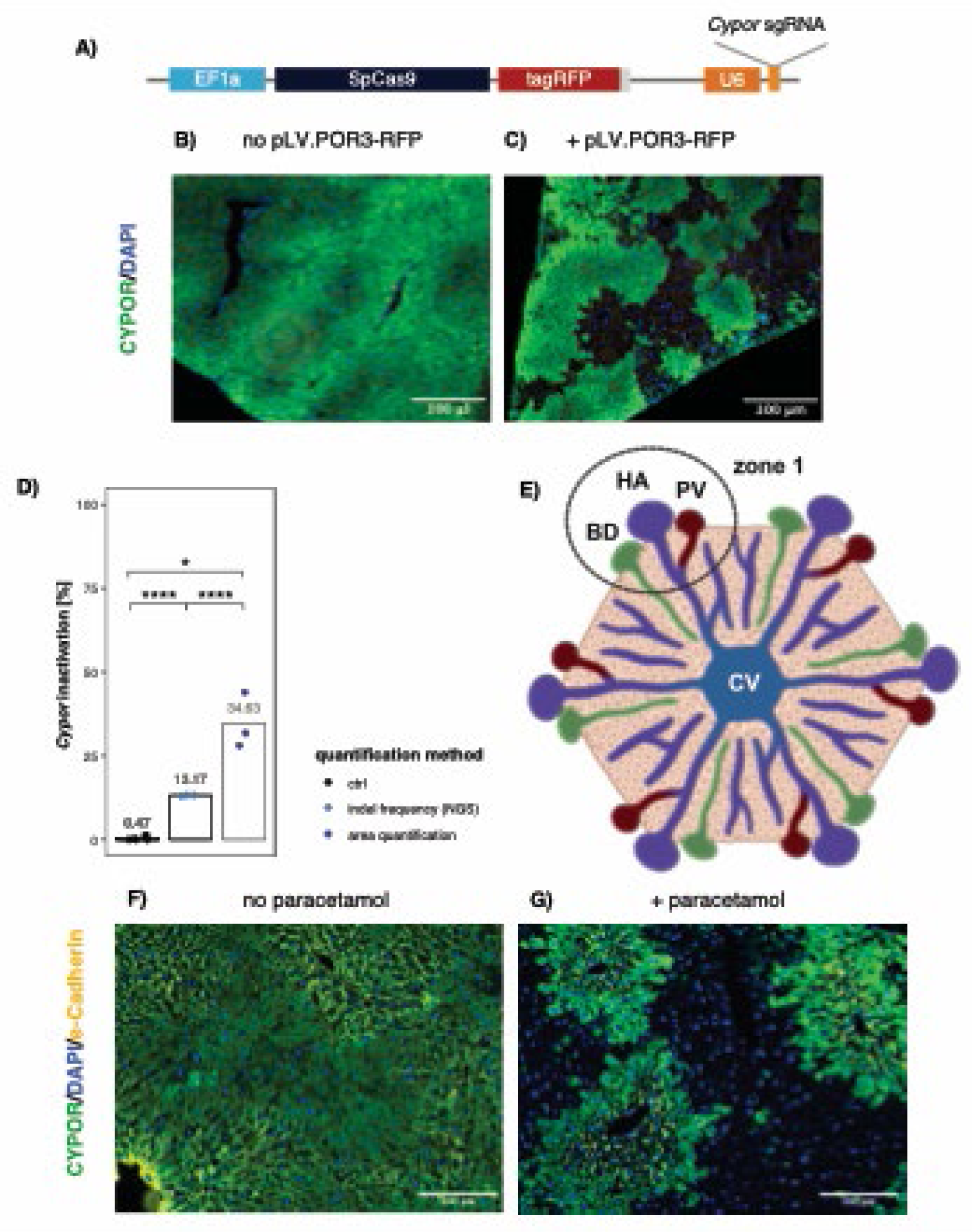
Selective perivenous-liver cell replacement upon Cas9-induced CYPOR inactivation in wild-type mice. (**A**) Schematic of the lentiviral vector plasmid pLV.POR3-RFP encoding SpCas9-tagRFP under the control of the ubiquitous EF1a promoter, and anti-Cypor gRNA controlled by the U6 promoter. (**B**) Liver staining of CYPOR expression in control wild-type mice treated with 13 paracetamol doses (no pLV.POR3-RFP vector administration; n = 2 mice). (**C**) Liver staining of CYPOR in wild-type mice (n = 3) treated with pLV.POR3-RFP vector, administered via HTV, and selected with 15 paracetamol doses (Scale bars, 100 μm; n = 3 mice). (**D**) Percentage of Cypor-inactivated gene copies detected using next generation sequencing (NGS), or microscopy areal quantification. (**E**) Schematic representation of a liver lobule, i.e. a collection of hepatocytes in a hexagonal shape with the center being a central vein (CV) in zone 3 while zone 1 contains the portal triad with bile duct (BD), hepatic artery (HA), and the portal vein (PV). (**F-G**) Representative liver staining for CYPOR and e-Cadherine of pLV.POR3-RFP vector (HTV) treated mouse, without paracetamol selection (**F**), and after 20 paracetamol doses (**G**; scale bars 200 μm. Green = CYPOR, yellow = e-Cadherin (periportal), blue = nucleus (DAPI). Statistics: unpaired t-test compared to ctrl and paired t-test comparing both quantification methods, * p < 0.05, ** p < 0.01, *** p < 0.001, **** p < 0.0001). E-Cadherin is specifically localized in the periportal-zone where hepatocytes are not replaced by paracetamol treatment as they do not express CYPOR. Note that although we used a LV vector expressing CYPOR-RFP, anti-RFP AB were not used for liver staining due to the strong red auto-fluorescence of liver tissue.

### Testing of lentiviral vector transduction and *Cypor*-gene editing efficiencies in Hep1-6 cells

While lentivirus (LV) is thought to be the most efficient vector to transduce primary hepatocytes (PH), transduction efficiency may highly vary between preparations^32^. We thus investigated transduction efficacy of LV.POR3-RFP (Fig. 1A) in the murine hepatoma cells line Hepa1-6 by determining the insertion frequency and the cutting efficiency, i.e. inactivation of *Cypor* by the *Sp*Cas9 and the corresponding anti-*Cypor* gRNA (see Methods section for selection of the gRNA). During the 4 hours incubation period, cells were supplemented with vitamin E and polybrene. As depicted in Fig. 2, three separate lots of LV preparations with MOI of 1’000, 2’000, and 5’000 vg/cell were tested and revealed on average of 2.1, 2.8, and 5.3 insertions per diploid genome and a corresponding cutting frequency of 52%, 76%, and 90%, respectively. Thus, neither transgene insertion nor its cutting frequency increase linearly with the viral titer. We thus decided to use 2’000 vg/cell for *ex vivo* primary hepatocyte transduction to limit the number of transgene insertion into the genome but also provide sufficient *Cypor* inactivation for successful *in vivo* selection.

**Figure 2.**
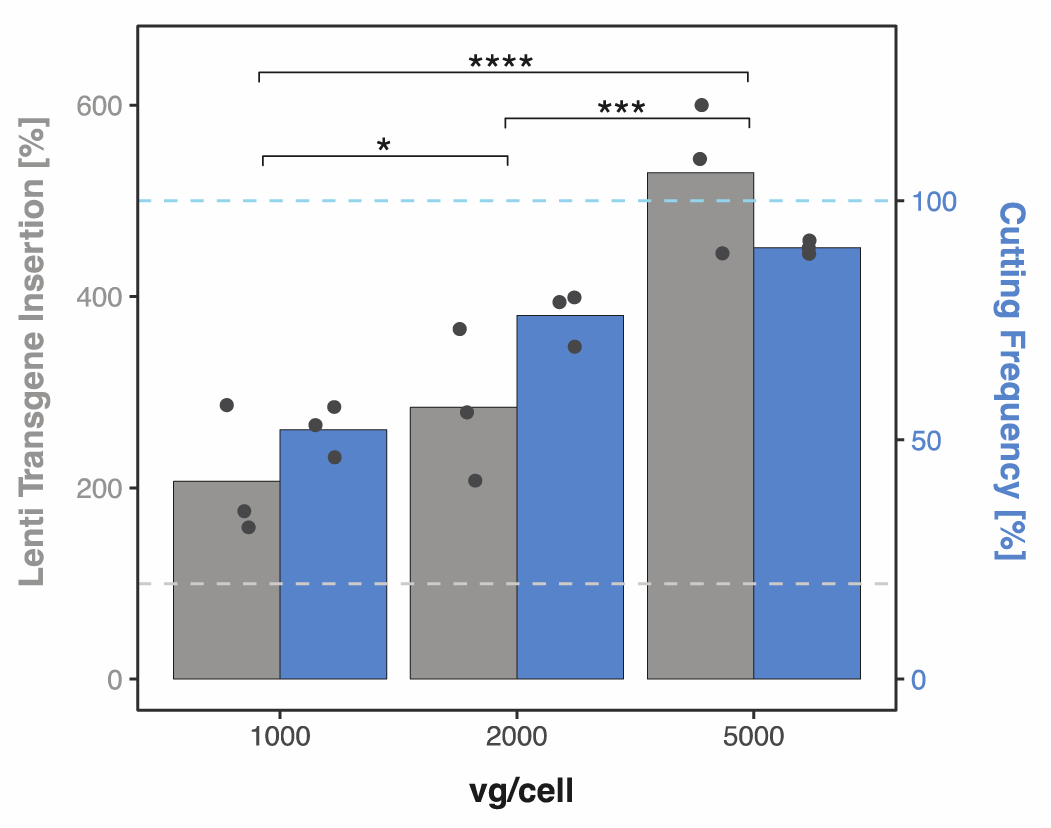
Validation of LV transduction using murine Hepa1-6 cells. Quantification of spCas9-tagRFP insertion and Cypor-inactivation of LV.POR3-RFP lots (n = 3) in Hepa1-6 cells transduced for 4 hours at 1000, 2000, and 5000 vg/cell. The lower and upper dashed lines represent 100% LV transgene insertion and 100% cutting frequency, respectively. Statistics: Two-way ANOVA, * p < 0.05, ** p < 0.01, *** p < 0.001, **** p < 0.0001).

### Successful treatment in murine PKU following in-vivo selection of transplanted hepatocytes

Based on our observations with pLV.POR3-FRP for the selective potential of paracetamol selection in wild-type mice (upon HTV infusion; Fig. 1) and the transduction efficacy of Hepa1-6 cells *in vitro* (Fig. 2), we applied and combined the procedures to treat a mouse model of PKU. Primary hepatocytes (PH) from wild-type mice that were isolated from inbred PKU mice and were incubated for 4 hours with LV.POR3-FRP (between 1’000 to 2’000 vg/cell; see below) for viral transduction and thereafter infused via the spleen into young adult PKU mice (5-8 weeks old, 4 males and 6 females; also see next section). Each mouse received 0.8 x 10^6^ PH, equivalent to approximately 0.5% of total liver hepatocytes^33^. While all male PKU mice, from three different transplantation cohorts, were grafted with 2’000 vg/cell-transduced donor wild-type PH. Female mice on the other hand were grafted with 1’000 vg/cell-transduced donor wild-type PH. The viral titer was reduced for female mice after the first cohort of females (n = 3) was grafted with 2’000 vg/cell-transduced wild-type PH, two out of three mice deteriorated rapidly within 24-48 hours post-transplantation and were terminated. Post transplantation, the transplanted mice underwent paracetamol selection by giving weekly moderate overdoses. Paracetamol selection was tightly followed in individual mice by weekly ALT measurements in combination with blood Phe levels (Fig. 3A, Fig. 5A, and Supplementary Fig. S2) monitored every 2-3 weeks. Three males (P65, P73, P99) and three females (P79, P105, P114) were treated successfully until Phe normalized (< 360 μM) and remained stable without selection thereafter for 2-4 additional weeks (Fig. 3A) with corresponding fur color change (Supplementary Fig. S3). Male mice required about 30 cycles of paracetamol for treatment whereas female mice required at least 40 paracetamol doses for treatment. Only the female PKU (P79) grafted with 2’000 vg/cell-transduced wild-type PH, reached normal blood Phe levels faster with only 33 paracetamol doses.

**Figure 3:**
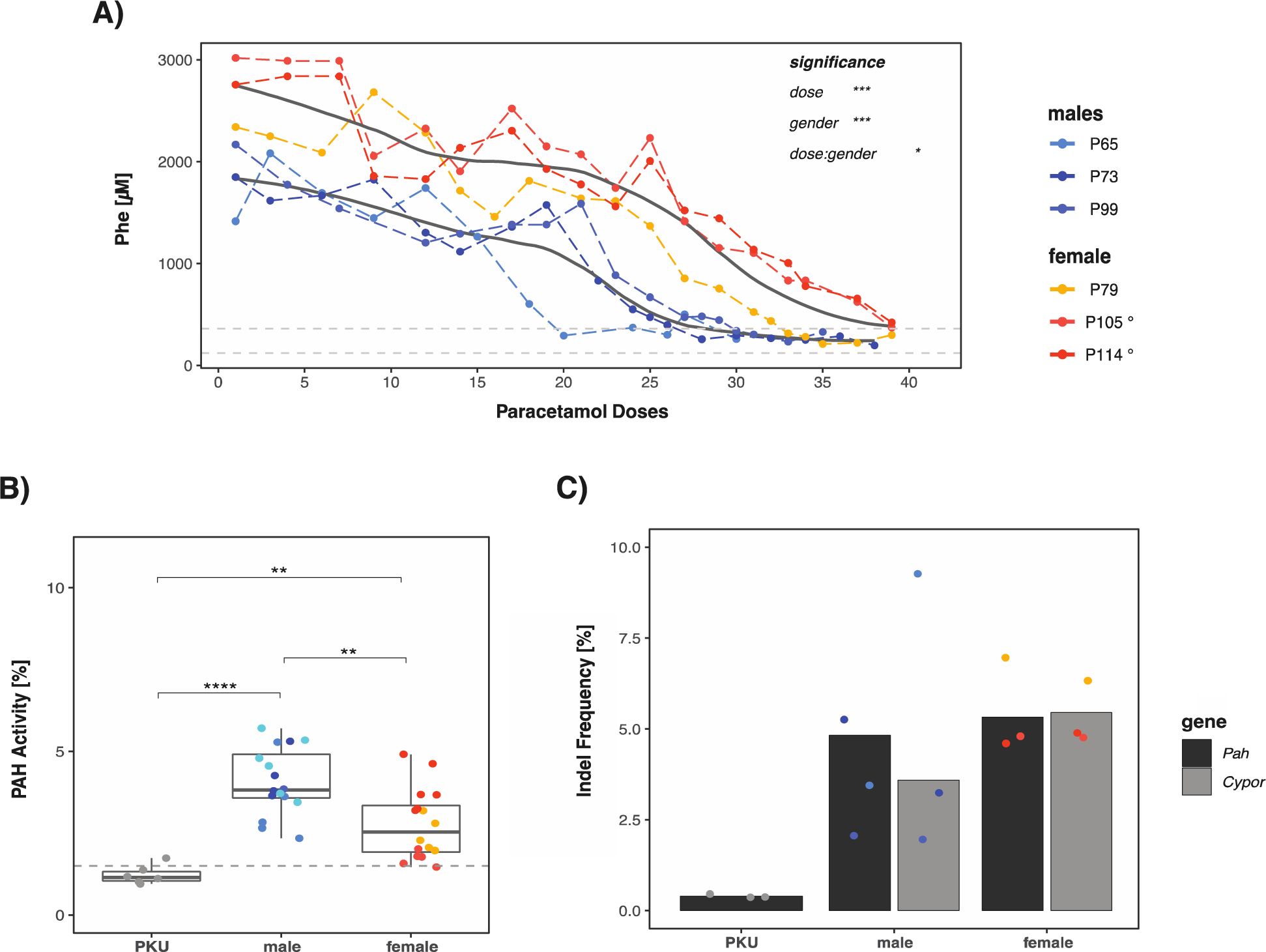
Successful treatment in murine PKU following ex-vivo gene therapy. (**A**) Change of blood Phe levels in male (n = 3) and female (n = 3) PKU mice during paracetamol dosing. The dashed lines represent the therapeutic threshold of 360 μM and “normal” blood Phe level which is below 120 μM, and the thick black lines represent the mean Phe level (Locally Weighted Scatterplot Smoothing using R). Note that one dose of paracetamol was given per week. At the end of the experiments, mice were sacrificed to determine PAH enzyme activity (**B**) and gene editing efficiency (**C**) in the resected livers. The % of PAH enzyme activity is presented relative to wild-type from all treated mice (the dashed line represents the PAH mean activity of PKU mice). Gene editing in individual mice was quantified by Cypor indel frequency determined by next generation sequencing (NGS). In addition, the percentage of wild-type Pah copy number was determined by NGS and all mice were transplanted with 2’000 vg/cell treated primary hepatocytes unless indicated otherwise (° = 1’000 vg/cell). Statistics: (A) Two-way ANOVA for unbalanced designs, (B) and (C) One-way ANOVA, * p < 0.05, ** p < 0.01, *** p < 0.001, **** p < 0.0001.

Treated PKU mice were monitored for blood Phe for an additional 2-4 weeks before euthanasia and liver resection. Post euthanasia, PAH enzyme activity and the percentage of transplanted, gene-modified wild-type hepatocytes were quantified from liver tissue. The mean PAH enzyme activity and wild-type *Pah* copy number was 3.5% ± 1.5% and 3.5% ± 1.6% for males and 2.2 % ± 1% and 5.5% ± 1.3% for females, respectively. The percentage of *Cypor* inactivation in the liver tissue was determined using next generation sequencing (NGS) and was 4.8% ± 3.9 % for males and 5.3% ± 0.9% for females, respectively. Additionally, IHC staining was performed on all mice where all five liver lobes were stained using anti-CYP2E1 and anti-CYPOR antibodies. As observed previously in mice treated via HTV with pLV.POR3-RFP and subsequent *in vivo* selection, the IHC staining confirmed selection of hepatocytes in the pericentral region of the liver (Supplementary Fig. S4). We also observed uneven engraftment and repopulation throughout the different liver lobes, which prohibited confident quantification using aerial quantification.

### Improvement of brain amino acid and neurotransmitter concentration

While systemic elevated Phe level does not affect the liver or lead to any liver pathology, it causes brain dysfunction due to perturbance of monoamine neurotransmitter metabolism with dopamine and (profound) serotonin deficiency in the central nervous system (by a still incompletely understood mechanism)^18^. We thus quantified the aromatic amino acids Phe, tyrosine (Tyr), and tryptophan (Trp), and the monoamine neurotransmitter metabolites dopamine, homovanillic acid (HVA), norepinephrine, serotonin, and 5-hydroxyindoleacetic acid (5-HIAA) in whole brain extracts of treated mice and controls. As expected, brain Phe, Tyr and Trp levels improved significantly but not completely in all treated PKU mice relative to untreated controls (Fig. 4A). Furthermore, serotonin was close to normal upon treatment, while the monoamine neurotransmitter metabolites 5-HIAA and norepinephrine improved to some extent compared to controls (Fig. 4B). Dopamine and HVA did not show any difference between PKU, treated PKU and wild-type mice. Overall, this brain metabolite analysis for PKU reflected what was known from similar studies with mice upon genetic, replacement or dietary treatment^34^.

**Fig. 4:**
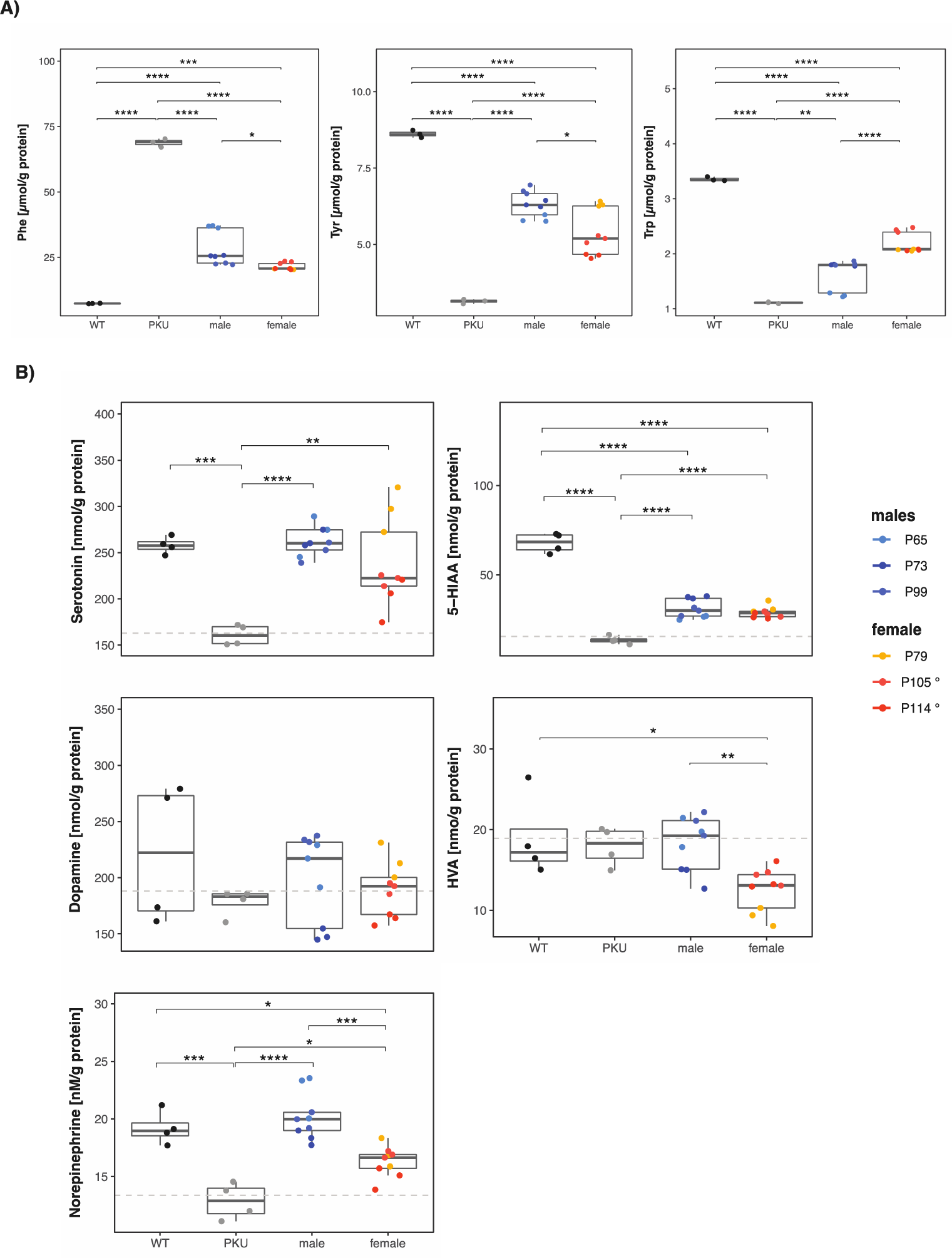
Brain amino acid and neurotransmitter metabolites in treated PKU mice. (**A**) Phe, Tyr, and Trp amino acid concentration in wild-type (WT) and PKU controls and male and female treated mice. (**B**) serotonin, 5-hydroxyindoleacetic acid (5-HIAA), dopamine, homovanillic acid (HVA), and norepinephrine neurotransmitter concentration in WT and PKU controls and male and female treated mice. Dashed lines represent mean concentration in PKU mouse brains. Statistics: Statistics: One-way ANOVA, * p < 0.05, ** p < 0.01, *** p < 0.001, **** p < 0.0001.

### Interrupting treatment delays and decreases Phe normalization

As previously mentioned, young adult PKU mice (5-8 weeks old, males n = 4, females n = 6) were grafted and selected *in vivo* for genetically modified hepatocytes, expressing wild-type PAH. The different cohorts (male-cohort 1: n = 1, cohort 2: n = 1, cohort 3: n = 2; and female-cohort 1: n = 1 (+ 2 deceased), cohort 2: n = 5; shown in Supplementary Fig. 2B) were defined by separate wild-type hepatocyte donors and transplantation dates but were treated as biological replicates. Throughout the paracetamol selection we were forced to interrupt the selection process multiple times unexpectedly, resulting in interruptions at different time points for each cohort (see Supplementary Fig. 2B). While blood Phe levels normalized after 30 doses in 3 out of 4 male PKU mice, 1 male PKU mouse’s blood Phe level remained around >1’300 µM. For female PKU mice, 3 out of 6 female PKU mice’s blood Phe levels normalized after 40 doses while the remaining 3 female PKU mice had blood Phe levels between 1’400 and 900 μM (Fig. 5A). When comparing all cohorts’ and the time points the paracetamol selection was interrupted, two observations were made. On one hand, multiple and prolonged interruptions during the paracetamol selection resulted in increased amounts of paracetamol doses required to reach successful blood Phe level normalization. On the other hand, if the selection process was interrupted prior to the 10^th^ paracetamol dose the success of blood Phe level normalization decreased by half. Nonetheless, preventing early interruptions during *in vivo* selection yielded 100% treatment success for males and females.

**Figure 5:**
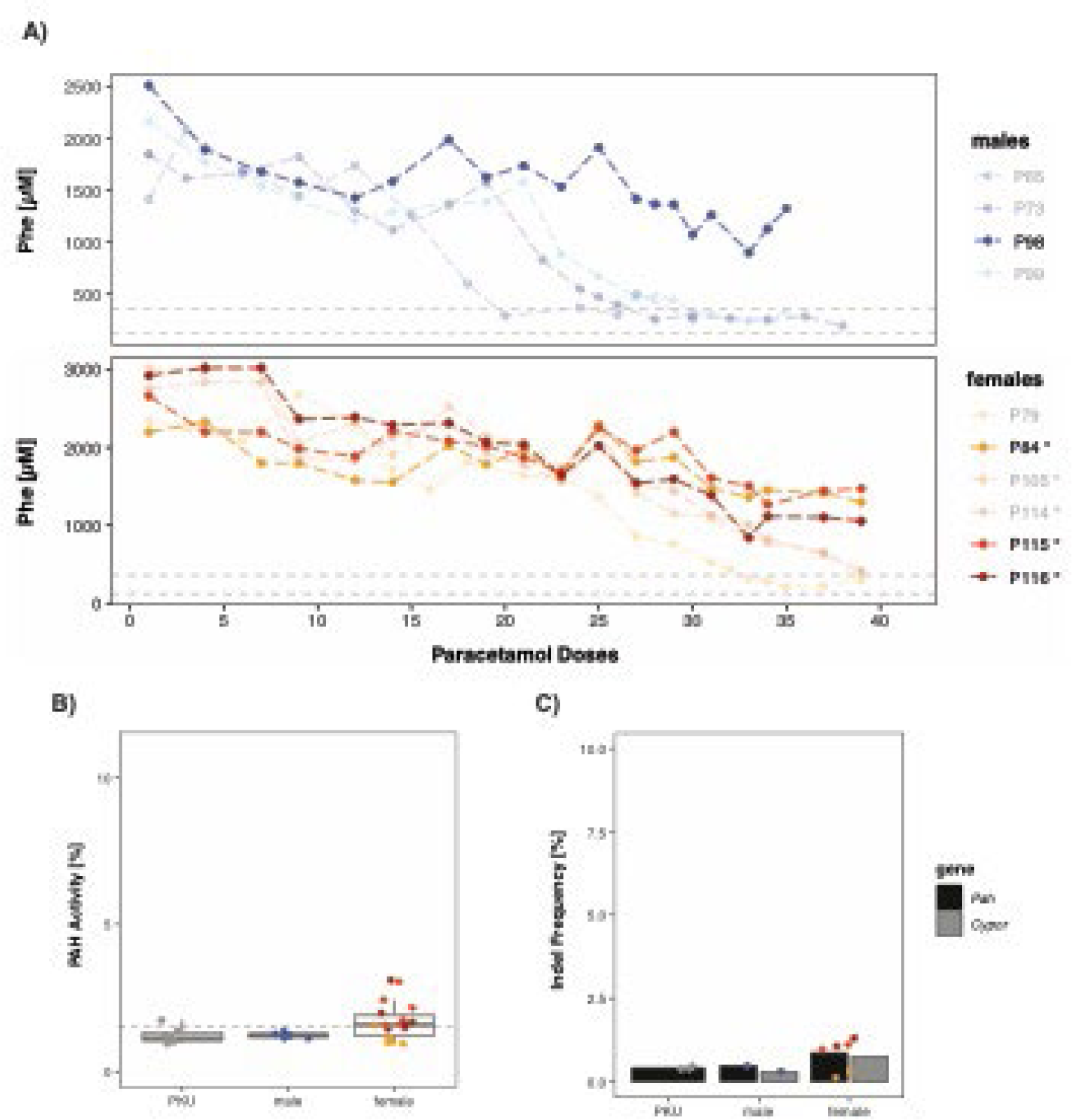
Paracetamol dosing for in vivo selection of modified hepatocytes following transplantation into PKU mice. (**A**) Change of blood Phe levels in male (n = 4) and female (n = 6) PKU mice during paracetamol dosing. The dashed lines represent the therapeutic threshold of 360 μM and “normal” blood Phe level which is below 120 μM, and the bold colored mice represent unsuccessfully treated mice. Note that one dose of paracetamol was given per week. At the end of the experiments, unsuccessfully treated mice were sacrificed to determine PAH enzyme activity (**B**) and gene editing efficiency (**C**) in the respected livers. The % of PAH enzyme activity is presented relative to wild-type from all treated mice (the dashed line represents the PAH mean activity of PKU mice). Gene editing in individual mice was quantified by Cypor indel frequency determined by next generation sequencing (NGS). In addition, the percentage of wild-type Pah copy number was determined by NGS and all mice were transplanted with 2’000 vg/cell treated primary hepatocytes unless indicated otherwise (° = 1’000 vg/cell). Note that interruptions were due to intervention by the animal welfare officer (AWO) of the State of Zurich for checks on animal welfare and compliance with the law.

## Discussion

Previous studies have shown that high DNA-vector doses delivered via HTV and viral gene transfer target 15-20% hepatocytes, sufficient to successfully treat IEM such as PKU ^20,30,31,35,36^. Nevertheless, when injecting naked DNA HTV or low viral vector doses encoding *Sp*Cas, it was shown that only 1-5% of hepatocytes show gene editing in adult mice^6,37^ rendering the editing efficiency at 10-25% in successfully targeted hepatocytes (Fig. 1). Nevertheless, using paracetamol-based *in vivo* selection the fraction of gene-edited hepatocytes can be increased from <2% to 13% with only 15 doses of paracetamol, increasing the fraction of gene-edited hepatocytes in the pericentral and midlobular region more than 6-fold. It remains an open question whether it will be feasible to increase the initially less than 1% gene-edited, *Cypor*-inactivated hepatocytes to reach levels of 15% or more since many delivery methods are less efficient than HTV or rely on high doses of viral vectors.

Monitoring of ALT response is a crucial component of *in vivo* paracetamol selection to monitor hepatoxicity and estimate degree of repopulation. During the paracetamol selection process of wild-type HTV-injected mice, we observed a significant increase in ALT response when the duration between doses was increased form 3-4 days to 7 days (Supplementary Fig. S1). We hypothesize that saturation of paracetamol metabolization and transient resistance to paracetamol overdosing, concomitant with liver hepatotoxicity may take up to 6 days^38^. Thus, within 7 days post paracetamol injection the liver can “regenerate” from overdosing and is able to metabolize paracetamol for the following selection round. We also observed gender associated differences where female PKU mice showed higher tolerance towards high paracetamol doses. Paracetamol selection dose to successfully decrease blood Phe levels increased form 230 mg/kg to 330 mg/kg and 280 mg/kg to >365 mg/kg for males and females, respectively. It has been shown that females are less susceptible to paracetamol-induced hepatotoxicity^39^ which requires further investigation for proper paracetamol dosage in potential patients. Additionally, we observed that the ALT response of each mouse within the same cohort may differ drastically (Supplementary Fig. S2) urging individual adjustment of paracetamol dosing.

LV vectors are highly efficient in transducing human PH *in vivo* and *ex vivo* but are substantially less efficient in transducing murine primary hepatocytes^40^. To assess the potential of LV.POR3-tagRFP for integration and *Cypor* inactivation in murine hepatocytes, we used Hepa1-6 cells for *in vitro* analysis. Here we observed that the insertion and cutting efficiencies did not exhibit a linear correlation (Fig. 2). Subsequently, we used 2’000 vg/cell for *ex vivo* PH transduction for which we observed 2-3 insertions per diploid genome and 75% *Cypor* inactivation in Hepa1-6 cells. Thereafter, grafting of up to 1 x 10^6^ PH per mouse was well tolerated which correlates to approx. 0.5% of total hepatocytes in the mouse liver. Assuming that all hepatocytes are engrafted, recipient mice must have less than 0.5% *Cypor*-inactivated wild-type hepatocytes in the liver upon transplantation. Eventually, liver-cell transplanted PKU mice normalized their blood Phe levels (below 360 μM) upon paracetamol cycling and without applying immune suppressants. While one male PKU mouse (P65) was treated with only 20 paracetamol doses, up to 30 paracetamol selection doses were required for the other PKU males (P73 and P99). Moreover, blood Phe remained at treatment level for up to three months without any further *in vivo* selection. Many gene therapy approaches for the liver have been applied successfully in males but not always in females. Here we observed that male PKU mice tolerated transplantation of 2’000 vg/cell-transduced PH. In contrast, female PKU mice (n = 2/3) showed severe complications within 24-48 hours upon transplanting 2’000 vg/cell-transduced PH, suggesting a gender specific immune reaction to the high LV titer used for *ex vivo* PH transduction; i.e., female PKU mice seemed to require a decreased lentiviral titer of 1’000 vg/cell for *ex vivo* PH transduction to tolerate subsequent PH transplantation. Only one female PKU was treated upon transplantation of 2’000 vg/cell-transduced PH and 33 paracetamol selection doses (P79; Fig. 3). By reducing the lentiviral load to 1’000 vg/cell for *ex vivo* PH transduction, all female PKU mice (n = 5) tolerated the transplantation well but required between 35-41 selection doses to reach therapeutic blood Phe levels. It remains an unexplained observation in humans (and many other mammals) that females show stronger immune responses to viral infections, including retroviruses, than males^41,42^.

During this *ex vivo* gene therapy experiment, we observed that the interruption during paracetamol selection (total of 10 weeks) directly correlated with increased paracetamol selection doses for successful treatment. Furthermore, one male and three females only showed decreased blood Phe levels but were not successfully treated (Supplementary Fig. S2). We conclude that interruption of *in vivo* selection, which realistically should be considered due to the possibility of unpredictable life events, may decrease therapeutic success by ∼50%. Especially at early time points during the *in vivo* selection, prior to 10^th^ paracetamol dose, fewer cells are *Cypor*-inactivated and interruptions should be avoided whereas interruptions during *in vivo* selection after the 10^th^ paracetamol dose will extend the total selection time but still allow successful repopulation. The decreased success rate may be explained by complications such as unwanted off-target transgene integration and the triggering of the adaptive immune system due to lentiviral chromosomal integration and stable transgene expression^43,44^. An additional limitation is introduced by the adaptive immune system which has been shown to induce preexisting humoral and cell-mediated immunity to the CRISPR-Cas9 system in many mammals^45-47^. Thus, replacing the lentiviral vector with a delivery vector allowing transient transgene expression may increase the treatment success rate despite interruptions at an early *in vivo* selection state.

In conclusion, we successfully treated male and female (*enu2*) PKU mice by a proof-of-concept *ex vivo* gene therapy approach using wild-type PH and subsequent *in vivo* selection. We could also show that even with (un-planned) interruptions during *in vivo* selection male and female PKU mice could be stably and successfully treated while the treatment was extended and the therapeutic success was decreased for some individuals. We believe the improvement and development of safer and more efficient transfer vectors for PH and the further understanding of paracetamol-induced selection provide the tools for future treatments for male and female patients affected by liver-based disorders.

## Animal handling and methods

### Mouse husbandry and handling

Animal experiments were approved by the State Veterinary Office of Zurich (Switzerland) and carried out according to the guidelines of the Swiss Law of Animal Protection (licenses ZH82-2019 and ZH118-2020), the Swiss Federal Act on Animal Protection (1978), and the Swiss Animal Protection Ordinance (1981). Mice were kept at controlled temperature, humidity, and a 12-hour dark-light cycle. Standard chow and drinking water were provided *ad libitum*.

### Hydrodynamic tail vein (HTV) injection

10 μg or 100 μg pLV.POR3-RFP plasmid was diluted in saline (0.9%, B. Braun Medical) to 10% mouse body weight. 8 week old male wild-type C57Bl/6J (Jackson Laboratory) or C57Bl/6J-PKU (*Pah^enu2/enu2^*) mice were anesthetized with 2-3% isoflurane in oxygen delivered with a flow rate 1 L/min and rapidly injected (5-8 seconds) with plasmid/saline solution via hydrodynamic tail vein (HTV) injection using a 27G needle^30^.

### Hepatocyte isolation, transduction and grafting

Primary hepatocytes (PH) were isolated from wild-type donor mice (same-sex) by a two-step collagenase digest. Briefly, mice were anesthetized using Ketamine (75 mg/kg), Acepromazine (2.5 mg/kg), and Xylazine (15 mg/kg) injected intraperitoneally (volume of 10 μL/g mouse). Upon full anesthesia, mice were perfused through the portal vein using a 24 G catheter with Ca2+/Mg2+-free Hanks’ solution (HBSS) containing 0.5 mM EGTA (for 5 min) and subsequently with Ca2+/Mg2+-containing HBSS with 0.1 mg/ml collagenase II (Worthington; for additional 15 min). All perfusion steps were performed using pre-warmed solutions (40°C) and a peristaltic pump (4 ml/min flow rate), ensuring a constant temperature and flow rate. Afterwards, the liver was returned to 4°C hepatocyte medium (high glucose Dulbecco’s modified Eagle’s medium (DMEM) substituted with 10% fetal bovine serum (FBS), 10 mM HEPES, 1% GlutaMAX, and 1% Penicillin/Streptomycin (Pen/Strep)), gently disrupted, filtered through a 100 μm, followed by 70 μm nylon cell strainer, and washed three times at 100 x g for 5 minutes to generate a single-cell, hepatocyte suspension. The cell yield was estimated using 0.04% Trypan Blue, where the viability ideally exceeded 80%.

For LV-transduction, PH were diluted to a concentration of 0.5 x 10^6^ cells/mL in hepatocyte media (described above) and transduced with 2,000 vg/cell for males, and 2,000 vg/cell or 1,000 vg/cell for females. To increase the transduction efficiency, 5 μL/ml of 10 nM vitamin E (in ethanol; Sigma Aldrich) and 1 μL/ml of 10 mg/ml polybrene (Sigma Aldrich) were added to the cell-virus suspension. 2 mL of cell-virus suspension was then seeded per 50 μg/mL rat tail collagen type I (Sigma Aldrich) coated 6-well plate wells. The cell-virus suspension was incubated for 4 hours and gently agitated every 20 minutes. After incubation the cells were transferred into a new 50 ml tube, cells that had attached were scraped off carefully, and washed with hepatocyte media three times at 100 x g for 5 minutes. The cell yield and viability were estimated using 0.04% Trypan Blue, where after the cells were diluted to a concentration of 8 x 10^6^ cells/mL, and stored at 4°C until transplantation.

For hepatocyte transplantation, 5-8 weeks old PKU mice were administered 10 mg/kg dexamethasone (Mephameson) 12 hours prior to transplantation to reduce a potentially adverse immunological response to LV. 30 minutes prior to surgery, mice received 1 dose of 5 mg/kg carprofen (Rimadyl, Pfizer) mixed with 0.1 mg/kg buprenorphine (Temgesic, Indivior Schweiz AG) for pain relief. Subsequently, the mice were anesthetized with 2-3% isoflurane in oxygen delivered with a flow rate 1 L/min, on a heating pad (37°C). The eyes were covered with Vitamin A, and the left flank was shaved and disinfected. The mice were transplanted via intra-splenic injection with 0.8 x 10^6^ viable, LV vector-transduced, primary wild-type hepatocytes in 100 μL hepatocyte medium using a 30G insulin syringe. After the procedure, 2 doses 5 mg/kg carprofen (Rimadyl, Pfizer) within 24 hours.

### Lentivirus (LV) production and analysis

On the LV expression vector pLV.POR3-RFP (13.3 kbp), *Streptococcus pyogenes Cas9* (*SpCas9*) was linked via P2A to tagRFP and controlled by the EF1a promoter. Four anti-CYPOR guide RNA (gRNA) were chosen from the Mouse Brie CRISPR knockout library^48^ (sgRNA-POR1: 5’-TGGCTCCCAGACGGGAACCG -3’, sgRNA-POR2: 5’-ATGTCTCTAAACAATCTCGA-3’, sgRNA-POR3: 5’-AAGAGGATTTCATCACATGG-3’, sgRNA-POR4: 5’-TCCAAGACTACCCGTCCCTG-3’). First gRNAs were cloned into a Lenti-SpCas9-Puromycin-gRNA plasmid (SpCas9-gRNA) using BsmBI, one gRNA per plasmid, and then tested for their cutting efficiency in Hepa1-6 cell line. Mouse Hepa1-6 cells were seeded on a 24 well plate at 1.5x10^5^ cells per well. One day after plating, cells were transfected by Lipofectamine 20000 (Thermo Fischer Scientific) according to the manufacturer’s instructions, with 600 ng of SpCas9-gRNA plasmid and 100 ng of a GFP plasmid as a transfection condition. For control conditions wild type Hepa1-6 cells and cells transfected with SpCas9-gRNA plasmid without gRNA sequence were used. Transfected cells were selected with puromycin (2.5 µg/mL) for 3 days. Nine days after transfection cells were harvested for DNA extraction using TrypLE-E (Gibco, 12605010). To extract DNA 100 ul of DirectPCR Lysis Reagent (Viagen) and 5 ul of Proteinase K were added to the cell pellet and incubated at 55 °C for 1 h. Proteinase K was inactivated at 85 °C for 10 min. To determine the gRNA cutting efficiency for the mouse *Cypor* locus DNA sequence around the Cas9 cutting site was analyzed. Briefly, genomic DNA was amplified by a PCR reaction using the Q5 Polymerase (NEB). Primer sequences are listed below (CYPOR-forward 5’-ACCCTCTCCTCTCCTCTCTC-3’ and -reverse primers 5’-TCCCCATCCTATGTCTGAGC-3’). PCR products were cleaned with magnetic beads (Sera-Mag Select, GE Life Science) and sent for Sanger sequencing using the CYPOR-reverse PCR primer. To quantify the gRNA cutting efficiency sequencing results were analyzed using the Tide software^49^. All experiments were performed in technical triplicates.

Vector pLV.POR3-RFP contained sgRNA-POR3 under the control of the U6 promoter, which was inserted using *BsmBI* restriction. The plasmids pMD2.G (Addgene 12259) and psPAX2 (Addgene 12260), were used to envelope and package. Briefly, four T150 tissue culture flasks of 80% confluent HEK293T cells in 15 ml complete media (high glucose Dulbecco’s modified Eagle’s medium (DMEM) substituted with 10% fetal bovine serum (FBS), 10 mM HEPES, 1% GlutaMAX, and 1% Penicillin/Streptomycin (Pen/Strep)) were triple-PEI-transfected at a molar ratio of 2:1:2 (pMD2.G:psPAX2:pLV.POR3-RFP) using a 1:3 DNA:PEI ratio in OptiMEM (Gibco). The medium was changed 20 hours post transfection and two batches were harvested 3 and 4 days post transfection. Supernatant was centrifuged at 450 x g for 5 minutes and subsequently filtered using a 0.45 μm filter into new tubes. Underlay the filtered supernatant with 10% v/v of 20% sucrose (in serum free DMEM + GlutaMAX) before centrifuging at 10,000 x g for 1.5 hours. Finally, the supernatant was discarded and the visible LV pellet was resuspended in remaining media, aliquoted and frozen on dry ice, and stored at -80°C.

### LV quantification using integration and cutting efficiency

Each LV:POR3-tagRFP (8.5 kbp) batch was tested for its integration and cutting efficiency as quality control. To determine the viral titer of a thawed LV aliquot, the Qubit7 RNA integrity and quality assay for quantification was used which is derived from the method used for AAV quantification^50^. Briefly, (i) a 5 μL sample was measured as is to determine the amount of free RNA, and (ii) a 5 μL sample was measured after 5 minutes of capsid denaturation at 95°C to measure the sum of free and encapsulated RNA. To obtain the amount of encapsulated RNA, the amount of free RNA was deducted from the total RNA. Thereafter, 0.6 x 10^6^ Hepa1-6 cells were transduced at 1,000, 2,000, and 5’000 vector genome (vg)/cell in 6-well plate with 5 μL/ml of 40 nM vitamin E (in ethanol; Sigma Aldrich) and 1 μL/ml of 10 mg/ml polybrene (Sigma Aldrich) and incubated for 4 hours. After 4 hours the cells were washed 2x and incubated for 7 days, during which each well was split into a 6 cm dish. At day 7, cell pellets were collected for DNA extraction which was extracted according to the manufacturer’s recommendation using DNeasy Blood & Tissue Kit (Qiagen, Hilden, Germany). To determine the integration frequency, a qPCR of tagRFP was performed using the primer set Frd: 5’-TCTGCAACTTCAAGACCACATA-3’, Rev: 5’-TCGGCCTCCTTGATTCTTTC-3’, probe: 5’-FAM-ACCCGCTAAGAACCTCAAGATGCC-TAMRA-3’ where tagRFP copy numbers were normalized to GAPDH (Mouse GAPDH Endogenous Control, Thermo Fisher). The cutting efficiency was quantified by amplifying the extracted gDNA using primer Frd: 5’-AGCCAGAGACTTCCAGTGCA-3’ and Rev: 5’-CCACACCTCACTGTGTCCCA-3’, cleaned up with NucleoSpin Gel and PCR clean-up (Macherey-Nagel), analyzed by Sanger sequencing, followed by Tracking of Indels by DEcomposition (TIDE) (version 3.3.0) analysis for indel frequency calculation.

### *In vivo* hepatocyte selection and liver repopulation

One week post-transplantation, paracetamol selection or cycling was started. Once per week, mice were fasted overnight for 16 hours to deplete consumed and stored glutathione. The following morning, mice were weighed, injected intraperitoneally with paracetamol (13 mg/ml in saline, ACROS Organics) at an initial dose for males and females of 230 mg/kg and 280 mg/kg, respectively. Thereafter, mice received standard chow *ad libitum*. 6 hours post paracetamol administration, they were weighed again and blood was collected from the tail vein for ALT measurement. ALT was analyzed as indicated by the manufacturer (Abbott Alinity C System, Abbot Laboratories, Chicago, IL, USA). In the following, mice were injected a paracetamol dose once per week for up to 41 weeks or until blood Phe-levels dropped below 360 μM, the defined target for hyperphenylalaninemia treatment^18^. The administered paracetamol dose was adjusted every week based on the prior week’s ALT level. To sustain a selective pressure, ALT response after paracetamol dosing must exceed 800 U/l, where the paracetamol was increased by 10 mg/kg when the ALT response was ≤800 U/l or 5 mg/kg when ≤2,000 U/l. If ALT level exceeded 2,000 U/l, the paracetamol dose was kept the same, or decreased by 5 mg/kg, if ALT level exceeded 4,000 U/l.

Blood Phe levels were determined approximately every second week by collecting dried blood spots on filter cards (after 3-4 hours of food fasting). Phe concentration was quantified by the Newborn Screening lab of the University Children’s Hospital by using isotope-dilution liquid chromatography-electrospray ionization tandem mass spectrometry (LC-ESI-MS/MS, Waters ACQUITY).

### Liver analysis

Upon euthanization of mice and saline perfusion, each liver lobe was sectioned into 3-5 mm thick sections and the rest was immediately frozen in liquid nitrogen. The sections were then fixed in 4% paraformaldehyde (PFA) at 4°C overnight where the remaining tissue was stored at -80°C. For genetic, enzymatic, amino acid, and neurotransmitter analysis the frozen liver and brain tissue were mechanically pulverized to powder using a Covaris tissue pulverizer (Covaris) to provide homogeneous analysis throughout the different liver lobes and the brain respectively.

### Indel frequency

gDNA was extracted from liver tissue powder using DNeasy blood and tissue kit (QIAGEN AG) according to the manufacturer’s instruction. For deep amplicon sequencing locus-specific primers (*oRT* − 255_*On* target 1F PKU: 5’-CTTTCCCTACACGACGCTCTTCCGATCTCCGTCCTGTTGCTGGCTTAC-3’, *oRT* − 256_*On* target 1R PKU: 5’-GGAGTTCAGACGTGTGCTCTTCCGATCTTGAGCATCCATTGTGGTTGG-3’, *oRT* − 824_*CYPOR*_*NGS*_*FW*: 5’-CTTTCCCTACACGACG CTCTTCCGATCTGAGTTGGGCCTTGGTGATG A-3’, and *oRT* − 825_*CY POR*_*NGS*_*Re*v: 5’-GGAGTTCAGACGTGTGCTCTTCCGATCTCTTCCACCCCGAAGAACTCG-3’) were used to generate targeted amplicons for deep sequencing. First, input genomic DNA was amplified in a 10 μL reaction for 26 cycles using NEBNext High-Fidelity 2x PCR Master Mix (NEB). PCR products were purified using Sera-Mag magnetic beads (Cytiva) and subsequently amplified for six cycles using primers with sequencing adapters. Approximately equal amounts of PCR products from each sample were pooled, gel purified, and quantified using a Qubit 3.0 fluorometer and the dsDNA HS Assay Kit (Thermo Fisher Scientific). Paired-end sequencing of purified libraries was performed on an Illumina MiSeq.

### PAH enzyme activity

Frozen liver tissues were lysed at 4°C in homogenization buffer adapted from^51^ using a QIAGEN TissueLyser device (QIAGEN AG). The phenylalanine hydroxylase (PAH) enzyme assay was then performed according to the assay developed in our lab previously^52^. Tyrosine concentration was determined by pre-column 6-aminoquinolyl-N-hydroxy-succinimidyl carbamate (AQC, Waters AccQ TagTM derivatization system) derivatization of the PAH enzyme assay product and subsequent separation by ultra-high performance liquid chromatography and UV absorbance detection (Waters AcquityTM UPLC, Milford, MA) using the Waters MassTrak Amino Acid Analysis method.

### Brain neurotransmitter analysis

Brain powder was homogenized, processed, and neurotransmitters were analyzed according to a previously published method^34^.

### Immunohistochemistry (IHC)

PFA fixed liver sections were moved to 30% (w/v) in phosphate buffered saline (PBS) until they sink to the bottom of the container. All liver tissue was then cryo-embedded, sectioned into 7 μm thick slices, dried, washed in PBS, and blocked in 2% normal donkey serum (Abcam) and 0.1% Triton X-100 in PBS for 30 min. Slides were incubated with primary antibodies rabbit anti-CYPOR (Abcam) (1:200) and goat anti-E cadherin (Abcam) (1:400) overnight at 4°C. 488-anti-rabbit (1:500) and Cy-anti-goat (1:500) were used as secondary antibodies, sections were counterstained with 4’,6-diamidino-2-phenylindole (DAPI), and mounted using Fluorescence Mounting Medium (Agilent). All images were taken using a Leica DMi8 confocal microscope.

### IHC, statistical analysis and graphical illustrations

Image analysis was conducted manually using ImageJ (version 2.9.0) where graphical illustrations (7.2 and 7.4e) were created with Biorender. All statistical analysis were performed and graphs were made using RStudio for macOS (version 2023.03.0.+386, Posit Software, PBC).

## Institutional Review Board Statement

Animal experiments were approved by the State Veterinary Office of Zurich (Switzerland) and carried out according to the guidelines of the Swiss Law of Animal Protection (licenses ZH82-2019 and ZH118-2020), the Swiss Federal Act on Animal Protection (1978), and the Swiss Animal Protection Ordinance (1981).

## Acknowledgments

This work was supported by grants (to B.T.) from the Swiss National Science Foundation (310030-162547 and Sinergia Grant CRSII5_180257), the Rare Disease Initiative Zürich (radiz) of the University of Zürich, and a mobility grant for doctoral students by the Swiss National Science Foundation to M.W. We thank the Newborn Screening unit for blood Phe measurements on dried blood spots, Dr. Claudia Lawnitzak (Veterinary Department of the Canton Zurich) for approving animal experiments, and Dr. Matthias Baumgartner (Division of Metabolism) for general support.

## Authors’ contribution

M.W., H.M.G.C., N.R., T.R., and M.H. performed experiments. M.W. and N.R. analyzed data. M.W, H.M.G., G.S, and B.T. conceived and managed the project. M.W. and B.T. wrote the manuscript with the contribution from all authors. All authors read and approved the manuscript.

