## Supplemental data for "Therapeutic liver cell transplantation to treat a genetic liver defect"

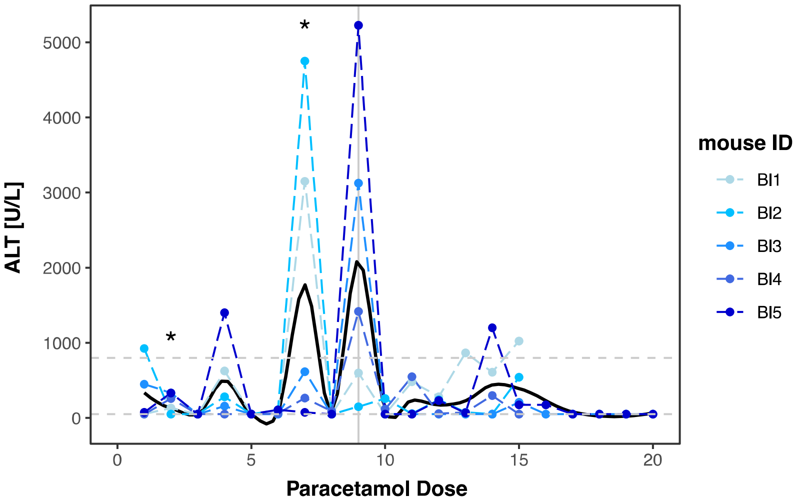


***Supplementary Figure 1. Alanine transaminase (ALT) monitoring during paracetamol dosing in wild-type mice upon HTV infusion.*** *ALT response was measured 6 hours post paracetamol administration in wild-type male mice (n = 5) infused with pLV.POR3-RFP. Mice were administered paracetamol every 3-4 days until dose 8, and only every 7th day from dose 9 on. The asterisk (*) marks paracetamol doses injected after 7 days prior to dose 9, and the vertical line marks paracetamol dose 9. Dashed horizontal lines represent the 50 U/l (lower line; background) and 800 U/l (upper line; selection threshold).*

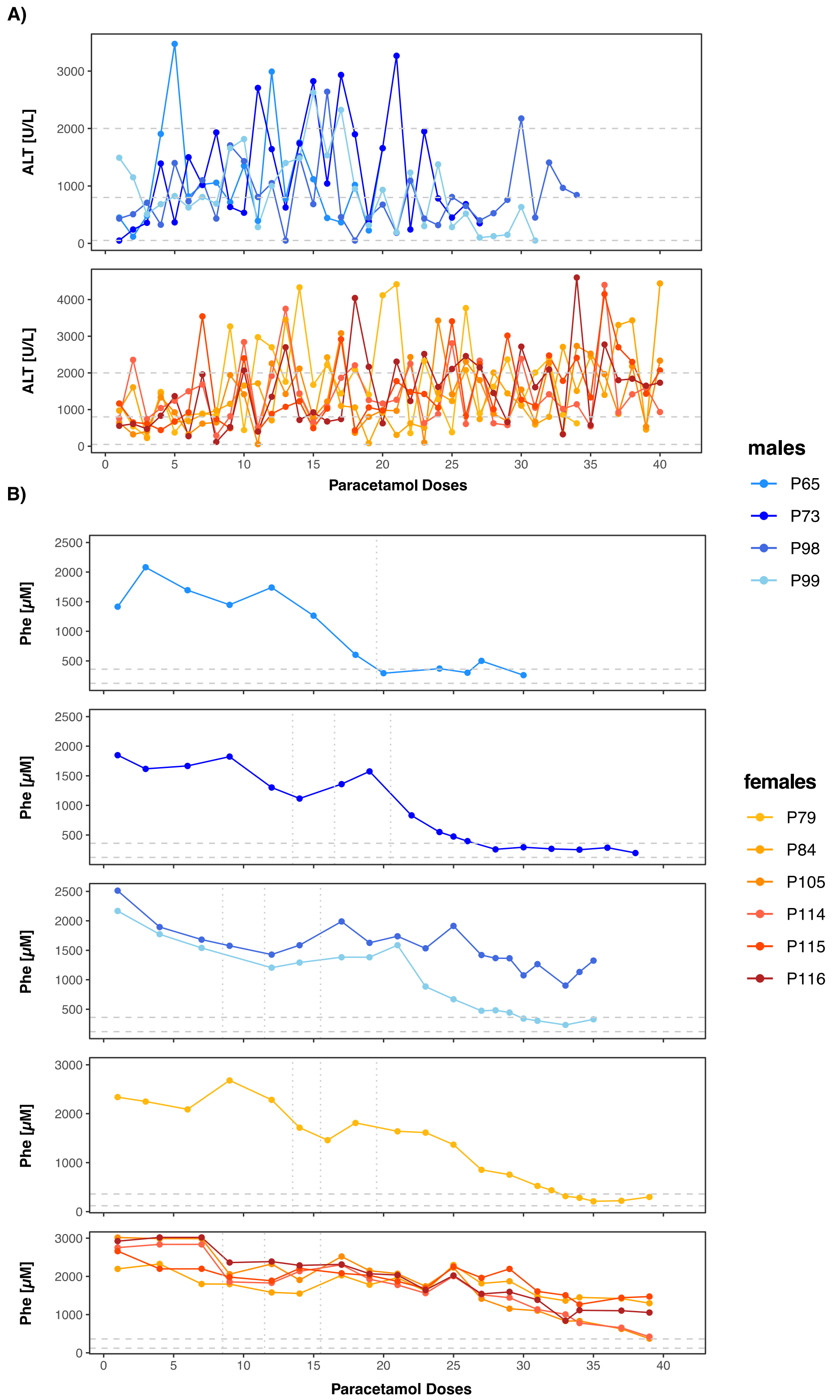


***Supplemental Fig. S2. Serum alanine transaminase (ALT) and blood Phe levels during paracetamol selection.*** *Young adult PKU mice (5-8 weeks old, males n = 4, females n = 6) were transplanted with wild-type hepatocytes, transduced by a selectable, Cypor-inactivating lentiviral vector, followed by paracetamol selection once per week. (**A**) ALT response in PKU mice during in vivo paracetamol selection. Dashed horizontal lines represent base line without selection at 50 U/L, minimal selection threshold at 800 U/L, and ideal upper limit for ALT response during selection. (**B**) Change of blood Phe levels in PKU mice during in vivo paracetamol selection split into corresponding cohorts. Dashed horizontal lines represent the therapeutic threshold of 360 µM and normal blood Phe level at 120 µM (blood withdrawal after 3-4 hours of feed deprivation). Note that the dotted vertical lines mark 2-4 week break from paracetamol selection.*

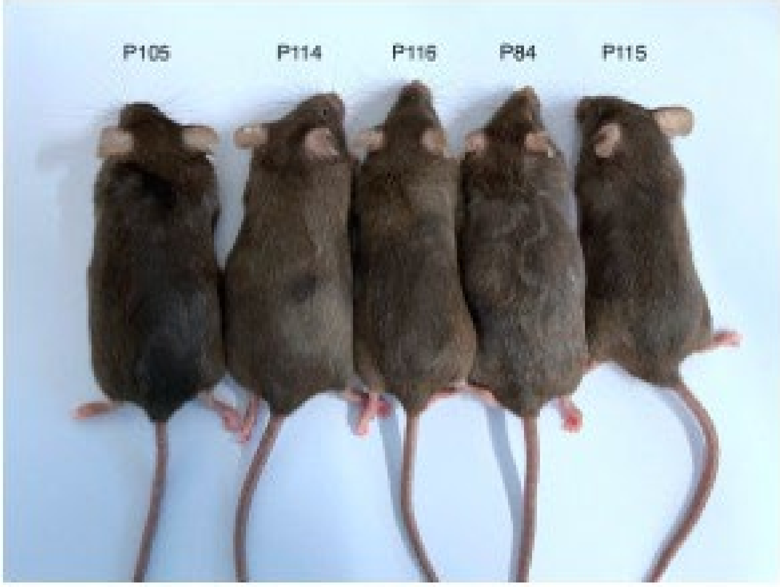


***Supplemental Fig. S3.*** *Examples of PKU female mice treated with 41 doses of paracetamol (for Phe levels see text and Fig. 3, Fig. 5, plus Suppl. Fig. 2).*

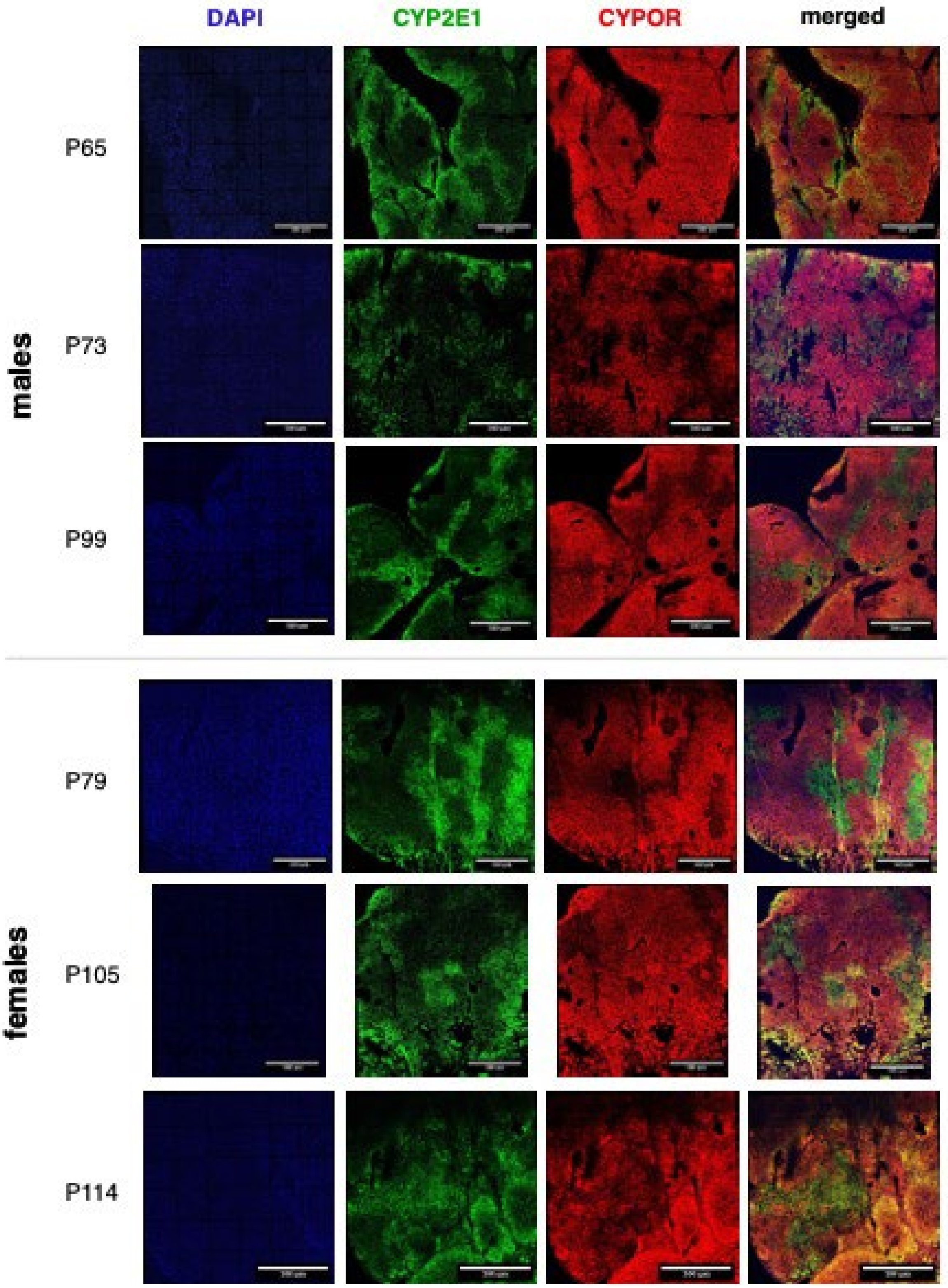


***Supplemental Fig. S4.*** *Examples of IHC staining of PKU male and female mice treated with up to 30 and 41 doses of paracetamol, respectively (for Phe levels see text and Fig. 3 plus Suppl. Fig. 2). From left, 1st panel (blue): nucleus (DAPI), 2nd panel (green): periportal (CYP2E1), 3rd panel (red): CYPOR, 4th panel: merge of panels 1-3. The pictures were taken at 10x magnification, and the scale bars is 500 µm.*
